# Multiple pathways to the evolution of positive assortment in aggregative multicellularity

**DOI:** 10.1101/2025.02.17.638078

**Authors:** Kayla S. Stoy, Kathryn A. MacGillivray, Anthony J. Burnetti, Cornelia Barrett, William C. Ratcliff

## Abstract

The evolutionary transition to multicellularity requires shifting the primary unit of selection from cells to multicellular collectives. How this occurs in aggregative organisms remains poorly understood. Clonal development provides a direct path to multicellular adaptation through genetic identity between cells, but aggregative organisms face a constraint: selection on collective-level traits cannot drive adaptation without positive genetic assortment. We leveraged experimental evolution of flocculating *Saccharomyces cerevisiae* to examine the evolution and role of genetic assortment in multicellular adaptation. After 840 generations of selection for rapid settling, 13 of 19 lineages evolved increased positive assortment relative to their ancestor. However, assortment provided no competitive advantage during settling selection, suggesting it arose as an indirect effect of selection on cell-level traits rather than through direct selection on collective-level properties. Genetic reconstruction experiments and protein structure modeling revealed two distinct pathways to assortment: kin recognition mediated by mutations in the *FLO1* adhesion gene and generally enhanced cellular adhesion that improved flocculation efficiency independent of partner genotype. The evolution of assortment without immediate adaptive benefit suggests that key innovations enabling multicellular adaptation may arise indirectly through cell-level selection. Our results demonstrate fundamental constraints on aggregative multicellularity and help explain why aggregative lineages have remained simple.

## Introduction

The evolution of multicellularity fundamentally altered the history of life. Evolving dozens of times independently (Grosberg and Strathmann 2007; Umen and Herron 2021), multicellularity facilitated cellular and morphological innovation, allowing multicellular organisms to radiate extensively across broad ecological niches (Stoy et al. 2024). Among these, ‘complex’ multicellular organisms have evolved a high degree of cellular specialization, morphological differentiation, and functional integration (Knoll 2011). Complex multicellularity has only evolved five times, occurring in animals, plants, red algae, green algae, and fungi (Knoll 2011; Brunet and King 2017). In contrast, all other multicellular organisms have remained comparatively simple, exhibiting little cellular and morphological specialization. Nascent multicellular lineages necessarily started out as simple collectives of cells (Bonner 1998; Pfeiffer and Bonhoeffer 2003; Grosberg and Strathmann 2007; Knoll 2011; Brunet and King 2017; Choi et al. 2024). In order to evolve into multicellular organisms, these collectives had to become Darwinian individuals: units of selection capable of gaining adaptations (Smith and Szathmary 1997; West et al. 2015; Godfrey-Smith 2019). In this paper, we examine a key constraint that inhibits this shift in the level of selection, from cells to collectives, for many multicellular organisms: the mechanisms by which natural selection acting on the traits of multicellular collectives can result in multicellular adaptation.

Multicellular development occurs through two basic modes: clonally and via aggregation. Clonal development occurs when individual cells remain permanently attached to one another following division, resulting in high genetic relatedness between cells (Bonner 1998). The evolution of complex multicellular life has only been observed in lineages that develop clonally (Knoll 2011; Brunet and King 2017), and there are several explanations for this macroevolutionary pattern. Clonal development limits opportunities for selection to act on cells within collectives because there is little within-group genetic variation upon which natural selection can act (Grosberg et al. 1998; Michod and Roze 2001; Fisher et al. 2013). Clonality can play an important role underpinning the heritability of novel multicellular traits by allowing selection on a multicellular trait (e.g., as an emergent property of mutations changing cell-level traits) to also act on the causal allele(s) shared by all cells in the collective (Herron et al. 2018; Bozdag et al. 2023; Zamani-Dahaj et al. 2023).

Alternatively, multicellular collectives can form though aggregation between previously free living and potentially genetically distinct cells, which can result in low within-group genetic relatedness (Bonner 1998; Velicer and Vos 2009; Tarnita et al. 2013; Du et al. 2015). This can constrain the ability of aggregative multicellular collectives to become Darwinian individuals (West et al. 2015; Godfrey-Smith 2019; Márquez-Zacarías et al. 2021). First, selection on cells within collectives can supersede selection on multicellular traits (Michod 1996; Michod and Roze 2001; Kuzdzal-Fick et al. 2011; Márquez-Zacarías et al. 2021). Second, within-group genetic diversity may result in a low covariance between genotype and multicellular phenotype, undermining the heritability of multicellular traits and blunting the efficacy of selection on these traits (Price 1970; Buss 1987; Michod 1997; Michod and Roze 1997; West et al. 2015; Godfrey-Smith 2019).

Aggregative multicellular organisms may overcome the challenge of low within-group relatedness by forming groups with positive genetic assortment, in which genetically similar cells preferentially adhere to one another. Many extant aggregative organisms have evolved kin recognition mechanisms (Ho et al. 2013; Bretl and Kirby 2016). However, little prior work has investigated whether such mechanisms create a covariance between multicellular phenotype and genotype that allows natural selection on collective-level traits to drive evolutionary change. In this study, we use aggregative strains of the yeast *Saccharomyces cerevisiae* to evaluate whether positive assortment facilitates multicellular adaptation in a nascent aggregative multicellular organism.

The yeast *S. cerevisiae* can form multicellular aggregates through a process called flocculation. “Flocs” form through lectin-like bonds between cell-surface FLO1 proteins and mannose oligosaccharides on adjacent yeast cell walls (Miki et al. 1982; Smukalla et al. 2008; Goossens et al. 2011; Brückner and Mösch 2012; Oppler et al. 2019). Cells expressing *FLO1* recognize and preferentially adhere to one another, excluding cells without *FLO1*, but flocs can be comprised of genetically diverse *FLO1*+ strains (Smukalla et al. 2008). We experimentally evolved twenty isogenic populations of flocculating *S. cerevisiae* under daily selection for both faster growth and the formation of large multicellular groups (Pentz et al. 2023). After 840 generations of settling selection for large multicellular size, flocs evolved to settle an average of 12-fold faster. Despite evolving increased settling rate, when flocculating yeast were competed against their ancestor, they exhibited no systematic fitness advantage (Pentz et al. 2023). We hypothesized this was due to the fast-settling evolved isolates formed chimeric groups with the slow-settling ancestor. This highlights a critical constraint for aggregative multicellular organisms: without a mechanism for generating positive assortment, selection on multicellular traits, even highly beneficial ones, cannot result in adaptive evolution.

In this study, we tested the hypothesis that positive assortment facilitates multicellular adaptation in a nascent aggregative multicellular organism. We predicted that if positive assortment facilitates multicellular adaptation, positively-assorted strains would gain a selective benefit. Competition assays between evolved aggregative *S. cerevisiae* strains and their ancestor revealed convergent evolution of positive assortment across 13 of 20 evolved replicate populations. Using several lineages that evolved mutations in the *FLO1* gene underlying cell-cell adhesion (Pentz et al. 2023), we then used genetic reconstruction experiments and protein folding simulations to assess potential underlying molecular mechanisms and evaluated whether recognition of partners at *FLO1* loci explained the observed positive assortment. Lastly, we evaluated potential phenotypic changes that may have increased positive assortment without requiring kin recognition. Together, these results demonstrate that positive assortment evolved without providing immediate adaptive benefits, suggesting it emerged indirectly through selection on cell-level traits that enhance flocculation efficiency rather than as a mechanism to enhance multicellular evolvability.

## Results

To test whether prolonged selection for large multicellular size favors the evolution of positive assortment in aggregative *S. cerevisiae*, we co-cultured evolved lineages with the ancestor and calculated assortment by measuring the relative frequency of evolved cells in flocs normalized by their overall frequency in the population, according to Eq. 1 (McNally et al. 2017):

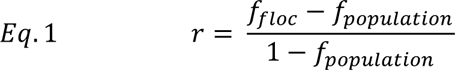

Assortment values can range from -1 to +1, with negative values indicating evolved cells are less likely to adhere to clonemates (negative assortment) than expected by chance, and positive values indicating preferential interactions with clonemates (positive assortment).

During experimental evolution, population F20 lost the ability to flocculate, so it was excluded from this analysis. We observed the evolution of increased positive assortment relative to the ancestor for 13 of 19 evolved lineages, and negative assortment for a single lineage (F9; Figure 1A, Table S1). Assortment values of these lineages were relatively low, with the highest assortment value of 0.328 and a mean value of 0.176 (Table S1). We then evaluated whether cells exhibiting positive assortment gained a fitness advantage during settling when competed against the ancestor during a single round of growth and settling selection (see methods in Pentz et al. 2023). We observed no significant relationship between assortment and the fitness of evolved lineages (Figure 1B; *F_1,17_* = 1.66, *R^2^* = 0.0357, *p* = 0.21). Together, these results suggest that evolved cells are more likely to join flocs, but preferential assortment itself is not adaptive during settling selection.

**Figure 1.**
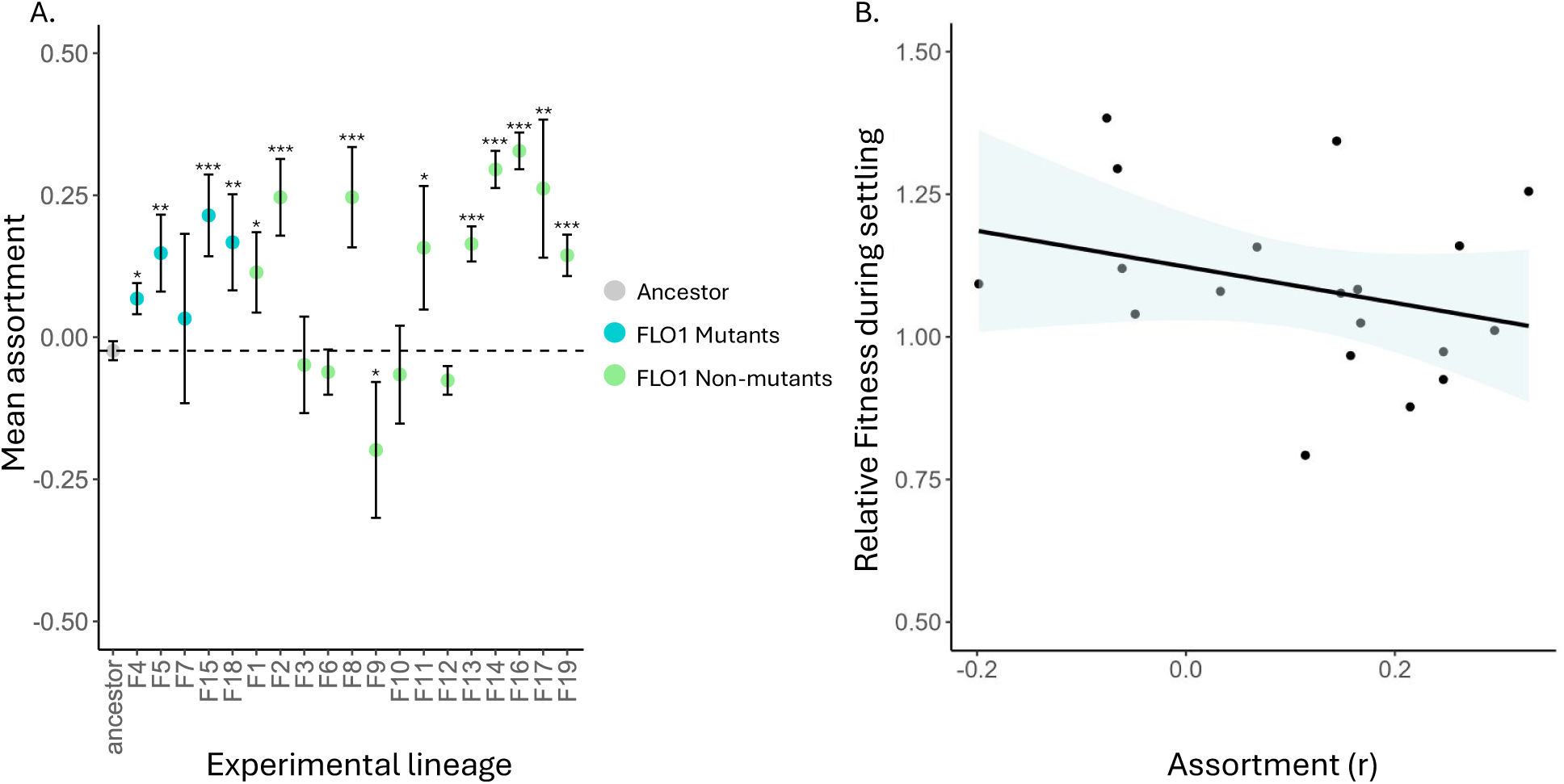
Assortment and settling advantage of evolved lineages. A) Mean assortment values for evolved lineages co-cultured with the ancestor. 13/19 lineages evolved increased positive assortment relative to the ancestor, while two lineages evolved negative assortment. *FLO1* mutants are lineages with evolved mutations at the *FLO1* gene. Dashed line represents the mean assortment value for differentially-labeled ancestral cells. Error bars show the standard error across independent assay replicates. Asterisks indicate a significant change from the ancestor. B) Linear regression of the assortment of evolved cells as a function of their selective settling advantage (measured as a selection rate constant (Lenksi *et al*., 1991) when competed against the ancestor. There is no significant relationship between assortment and selective settling advantage (*F_1,17_* = 1.66, *R^2^* = 0.036, *Pearson’s r* = -0.299, *p* = 0.21), indicating that evolving positive assortment alone does not confer a fitness benefit during competition with the ancestor. Blue shaded region denotes a 95% confidence interval.

The lack of a fitness benefit from positive assortment was surprising, particularly in the context of settling selection. These lineages evolved to settle 12-fold faster than their ancestor when measured in isolation, and the capacity to exclude their slow-settling ancestor from aggregative flocs should, in theory, provide considerable fitness advantages during settling selection. Specifically, cells settling in groups composed primarily of fast-settling individuals should outcompete those in mixed groups containing slow-settling ancestors. However, we observed no such advantage, suggesting that positive assortment was not under direct selection despite its potential benefits for multicellular adaptation. To better understand this puzzling observation, we investigated the potential mechanisms underlying positive assortment in our experimental system.

The *FLO1* gene mediates cell-cell adhesion within aggregates via binding to cell wall carbohydrates (Miki et al. 1982; Van Mulders et al. 2009; Goossens et al. 2011; Oppler et al. 2019), and cells expressing the *FLO1* gene recognize and preferentially adhere to one another (Smukalla et al. 2008). Four of the experimentally evolved lineages convergently evolved point mutations in the *FLO1* gene (Pentz et al. 2023), which may result in structural changes to the protein binding site of the FLO1 protein (see Table S2; Lineage F4 evolved a large deletion at *FLO1* and was excluded from analysis in this manuscript). To examine the potential consequences of these mutations on the FLO1 protein, we used simulations in AlphaFold 3 (Abramson et al. 2024). Including the evolved mutations S51F (lineage F15), N84D (Lineage F5), K194D (Lineage F7), and P201R (Lineage F18) in simulations did not predict gross changes in folding as a result of these *FLO1* mutations. However, the four independent mutations occur in a similar location: in and around the carbohydrate binding pocket of the N-terminal ligand binding domain of FLO1p (Figure 2).

**Figure 2.**
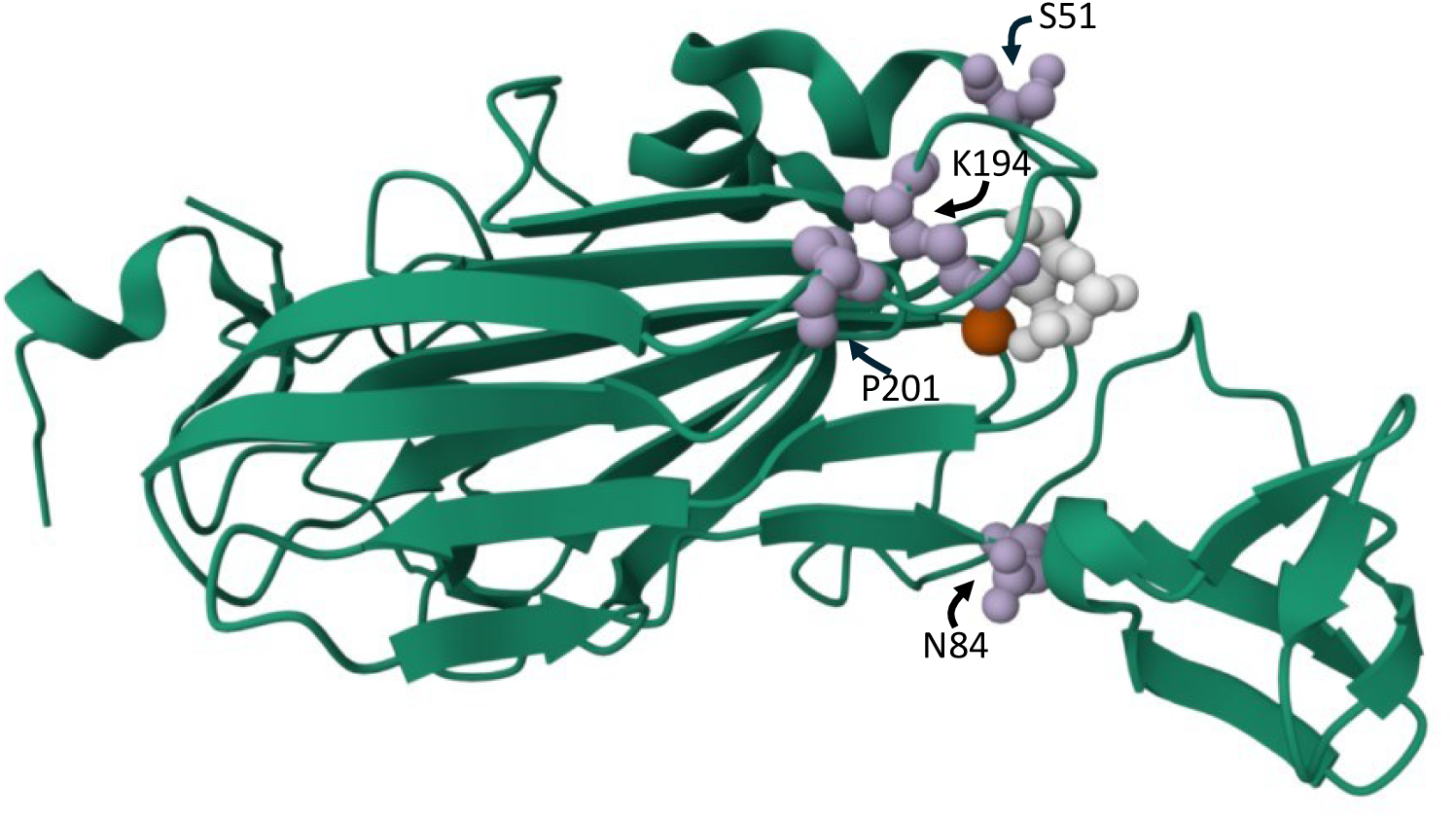
Visualization of simulation of the FLO1 protein with mutations. Locations of observed mutated residues (S51, N84, K194, P201) on structure of the FLO1 N-terminal carbohydrate binding domain [PDB structure 4LHN (Goossens et al. 2015; Lelasi and Willaert 2023)] in complex with calcium (red) and mannose (white). Residue K194 (lineage F7) directly contacts bound mannose via hydrogen bonds in crystal structures of FLO1, while others surround the edges of the primary binding site. P201 (lineage F18) defines the edge of the flexible “L3” loop participating in binding, N84 (lineage F5) is located at a ‘hinge’ to a subdomain also involved in binding, and S51 (lineage F15) while not part of the binding site is adjacent to it.

Because the FLO1 protein is known to play a role in cell-cell adhesion via binding to cell wall carbohydrates, and the observed mutations cluster around the carbohydrate binding pocket, we predicted these evolved changes to *FLO1* could underpin positive assortment and resulted in kin recognition, such that cells with the same FLO1 protein recognized and preferentially adhered to one another due to altered carbohydrate binding and cell wall composition. To test this, we replaced the ancestral allele with the evolved *FLO1* allele from each of the evolved *FLO1*-mutant lineages (hereafter referred to as the ANC+FLO1_evo_ lineages). ANC+FLO1_evo_ lineages were co-cultured with the ancestor, and we measured assortment, as described above. We observed an increase in positive assortment relative to the ancestor for the F5 ANC+FLO1_evo_ lineage (Figure 3A; *t* = -2.92, *df* = 6, *p* = 0.04), and the mean assortment value observed for F5 ANC+FLO1_evo_ (*r* = 0.120) was similar to the mean assortment value of the F5 lineage (*r* = 0.148) (Figure 3A). These observations suggest that changes to the F5 FLO1 protein underlie the observed patterns of positive assortment, which is consistent with kin recognition mediated by this allele.

**Figure 3.**
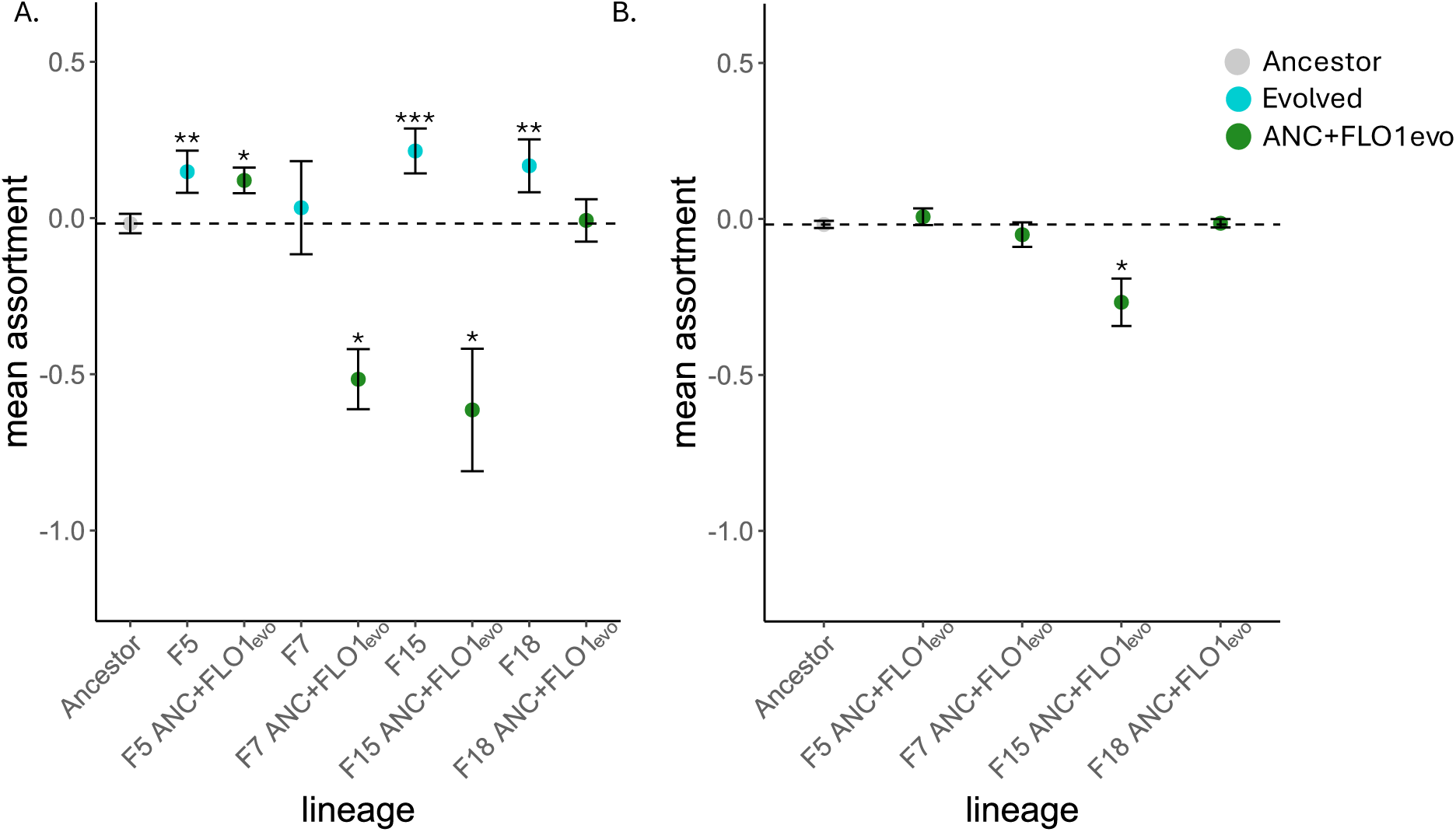
Assortment of engineered ANC+FLO_evo_ strains. A. Assortment of ANC+FLO_evo_ strains when co-cultured with the ancestor. Assortment of the evolved FLO1-mutants is included for visualization, and changes from the ancestor were analyzed separately (as described above). The F5 ANC+FLO_evo_ strain exhibited a significant increase in positive assortment relative to the ancestor. F7 ANC+FLO_evo_ and F15 ANC+FLO_evo_ exhibited a significant change from the ancestor toward negative assortment relative to the ancestor. The assortment of the F18 ANC+FLO_evo_ strain did not differ from that of the ancestor. B. Assortment of ANC+FLO_evo_ strains when co-cultured with evolved cells sharing the same *FLO1* gene. The engineered F5 ANC+FLO_evo_, F7 ANC+FLO_evo_, and F18 ANC+FLO_evo_ exhibited neither positive nor negative assortment, such that ANC+FLO_evo_ cells recognized and formed flocs with evolved cells that shared the same *FLO1* gene. The F15 ANC+FLO_evo_ lineage exhibited negative assortment, indicating an inability to recognize and adhere to cells from the evolved F15 that shared the same FLO1 gene. Dashed lines represent the mean assortment value for differentially GFP- and RFP-labeled ancestral cells. Mean assortment values are shown in grey for ancestor, blue for evolved cells, and green for the engineered ANC+FLO_evo_ strains. Error bars show the standard error in assortment values across independent assay replicates. Asterisks indicate a statistically significant change from the ancestor.

In contrast, the *FLO1* mutation was insufficient to explain the positive assortment observed in lineage F18 (Figure 3A, *t* = -0.54, *df* =6, *p* = 0.61). Surprisingly, the F7 ANC+FLO1_evo_ and F15 ANC+FLO1_evo_ strains exhibited a significant change from the ancestor toward negative assortment (Figure 3A; F7: *t* = 4.44, *df* = 6, *p* = 0.02; F15: *t* = 2.81, *df* = 6, *p* = 0.04). For these lineages, replacing the ancestor’s *FLO1* allele with an evolved allele fundamentally reduced this lineage’s ability to enter flocs, suggesting that *FLO1* in lines F7 and F15 evolved with other elements of the cell which FLO1p binds to, like the composition of the cell wall. Placing the *FLO1* gene from the evolved F7 and F15 lineages into the ancestral genomic background reduced their ability to flocculate. Therefore, changes to the *FLO1* gene alone could not explain the increased flocculation efficiency of these lineages nor the increased positive assortment observed for the F15 evolved lineage.

Given that cell-cell adhesion requires both FLO1-protein interactions and cell wall binding, we investigated whether isolating evolved *FLO1* alleles from their cellular context might disrupt their function. To test this possibility, we co-cultured each engineered ANC+FLO1_evo_ lineage with cells from the evolved lineage that shared the same *FLO1* gene. When F7 ANC+FLO1_evo_ was cultured with the F7 lineage, we observed a mean assortment value of *r* = -0.054, which did not significantly differ from the assortment observed for the ancestor (*r* = -0.0177; *t* = 0.80, *df* = 6, *p* = 0.61). These results suggest that flocculation efficiency depends on the broader cellular context in which the FLO1 protein functions, not just the *FLO1* allele alone. When F15 ANC+FLO1_evo_ was cultured with the F15 lineage, we observed a significant change from the ancestor toward negative assortment (*r* = -0.267; *t* = 3.84, *df* = 5, *p* = 0.049). The reduced capacity of the F15 ANC+FLO1_evo_ cells to join flocs may indicate that removing the F15 *FLO1* gene from the cellular context in which it evolved disrupted its function. We observed assortment values close to zero for the F5 ANC+FLO1_evo_ (*r* = 0.007) and F18 ANC+FLO1_evo_ (*r* = -0.014) lineages, which did not significantly differ from the assortment of the ancestor. This was consistent with expectations given these ANC+FLO1_evo_ lineages readily entered flocs when competed against the ancestor.

The analyses above suggest multiple evolutionary pathways resulted in the evolution of positive assortment. For F5, changes to *FLO1* appeared to mediate positive assortment. However, changes to *FLO1* alone were insufficient to result in positive assortment for the remaining *FLO1*-mutant lineages. Positive assortment was also observed for evolved lineages without mutations in *FLO1* (Figure 1A). Together, this suggested that cellular changes independent of *FLO1* could also result in positive assortment, potentially through modifications to cell adhesion or other aspects of cell-cell interactions.

Cells could achieve positive assortment through several mechanisms, including changes that make them generally more adhesive and better able to form strong bonds. For example, increased mannose production (which FLO1p binds to) could enhance cell-cell adhesion and provide an alternative pathway to positive assortment independent of *FLO1* mutations. Adding mannose to cell culture media can competitively inhibit binding between cells (Eddy 1955; Mill 1964; Miki et al. 1982). Therefore, we used competitive inhibition assays to test this prediction by adding mannose to cell culture media (YPG) at concentrations of 0%, 1%, 2.5%, 5%, and 7% and measuring flocculation efficiency of the evolved *FLO1*-mutant lineages and the ancestor.

To investigate how the flocculation efficiency of the evolved lineages differed from the ancestor in response to increasing mannose concentrations, we fit a linear regression with an interaction term between lineage and mannose concentration (treated as a continuous variable). Flocculation efficiencies of each evolved lineage were normalized by the ancestor. By comparing the slopes of the regression for each evolved lineage with that of the ancestor (*m* = 0), we assessed how the evolved lineages compared to the ancestor in their responses to increasing mannose concentrations. Lineage F5 did not differ from the ancestor in its response to mannose (*m* = -0.0162, *95% CI* [-0.0521, 0.0197]) (Figure 4A). However, the slopes of the regressions for lineages F7 (*m* = -0.0776, *95% CI* [-0.1135, -0.0417]), F15 (*m* = -0.0468, *95% CI* [-0.0827, -0.0109]), and F18 (*m* = -0.1243, *95% CI* [-0.162, -0.0884]) were significantly different from zero (Figure 4A), indicating they differed from the ancestor in their response to increasing mannose concentrations. These lineages are more sensitive to exogenous mannose than the ancestor, though this may be because they start out with much higher flocculation efficiency (F7 intercept = 0.749, *p* < 0.0001, F15 intercept = 0.39, *p* = 0.0003, F18 intercept = 0.96, *p* < 0.0001; Figure 4), and thus have further to fall when flocculation is disrupted.

**Figure 4.**
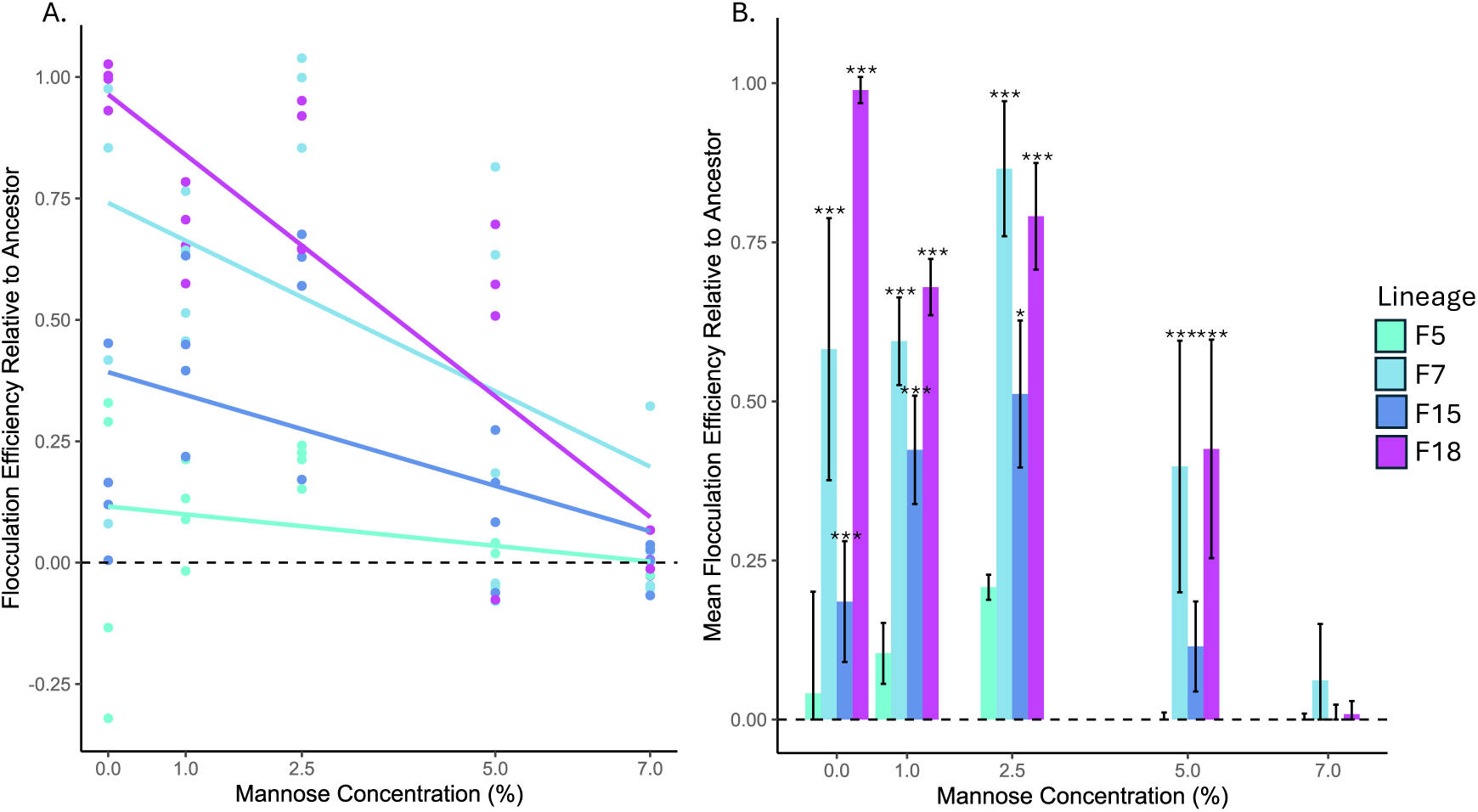
Competitive inhibition of flocculation by exogenous mannose. A. Linear regressions of flocculation efficiency as a function of mannose concentration for the evolved lineages. Flocculation efficiency values were normalized to the ancestor (dashed line). The slopes of the regressions for the evolved lineages F7, F15, and F18 are significantly different from zero, indicating these lineages differed from the ancestor in their response to increasing mannose concentrations. Points show the flocculation efficiencies of each lineage for each independent assay replicate. The dashed line shows the ancestor. B. Mean flocculation efficiency of evolved strains normalized by the ancestor at each mannose concentration. Lineages F7, F15, and F18 exhibited significantly higher flocculation efficiencies than the ancestor at 0%, 1%, and 2.5% mannose. Lineages F17 and F18 exhibited significantly higher flocculation than the ancestor at 5%. At 7% mannose, none of the evolved lineages differed from the ancestor. Dashed line shows the ancestor. Error bars show the standard error across independent assay replicates.

We then compared the flocculation efficiencies of the evolved lineages and the ancestor at each specific mannose concentration. To do this, we used estimated marginal means to predict pairwise relationships based on the linear regression. Lineages F7, F15, and F18 all exhibited higher flocculation efficiency than the ancestor at 1% (F7: *p* < 0.0001, F15: *p* = 0.001, F18: *p* < 0.0001) and 2.5% (F7: *p* < 0.0001, F15: *p* = 0.001, F18: *p* < 0.0001) mannose concentrations (Figure 4B). At 5% mannose, lineages F7 (*p* = 0.0004) and F18 (*p* = 0.0007) exhibited significantly higher flocculation efficiency than the ancestor (Figure 4B). At 7% mannose, flocculation was nearly inhibited for all lineages and none of the evolved lineages differed from the ancestor (Figure 4B). Overall, lineages F7, F15, and F18 maintained their ability to form aggregates at higher mannose concentrations compared to the ancestor, suggesting these evolved strains formed stronger cell-cell bonds that were more resistant to competitive inhibition by free mannose.

If positive assortment resulted from preferential binding to kin, we expected related cells would not only preferentially form flocs with one another, but would exhibit spatial structure within flocs. Alternatively, if positive assortment resulted from generalized increased adhesiveness (e.g., increased mannose production), we would not expect to see intrafloc spatial structure. To test for spatial structure within flocs, we co-cultured fluorescently labeled evolved and ancestral cells. We then flattened floc isolates into a two-dimensional distribution (Figure 5B) and calculated the frequency of interactions between evolved cells within a 4 µm radius (cell radius is approx. 2.5 µm). While some elements of the 3D structure of the floc are destroyed by flattening, short spatial interactions (*i.e.*, bonding between nearest neighbors) should largely be retained. Further, this approach is conservative, as this method should not systematically create assortment, though it may disrupt it. For each lineage, we simulated the random distribution of evolved and ancestral cells over 1000 iterations to create a null distribution of the frequency of interactions between evolved cells expected at random. We compared the null distribution for each lineage to the observed data and assessed the proportion of simulated interaction frequencies outside of the 95% confidence intervals of the observed data (Figure 5A).

**Figure 5.**
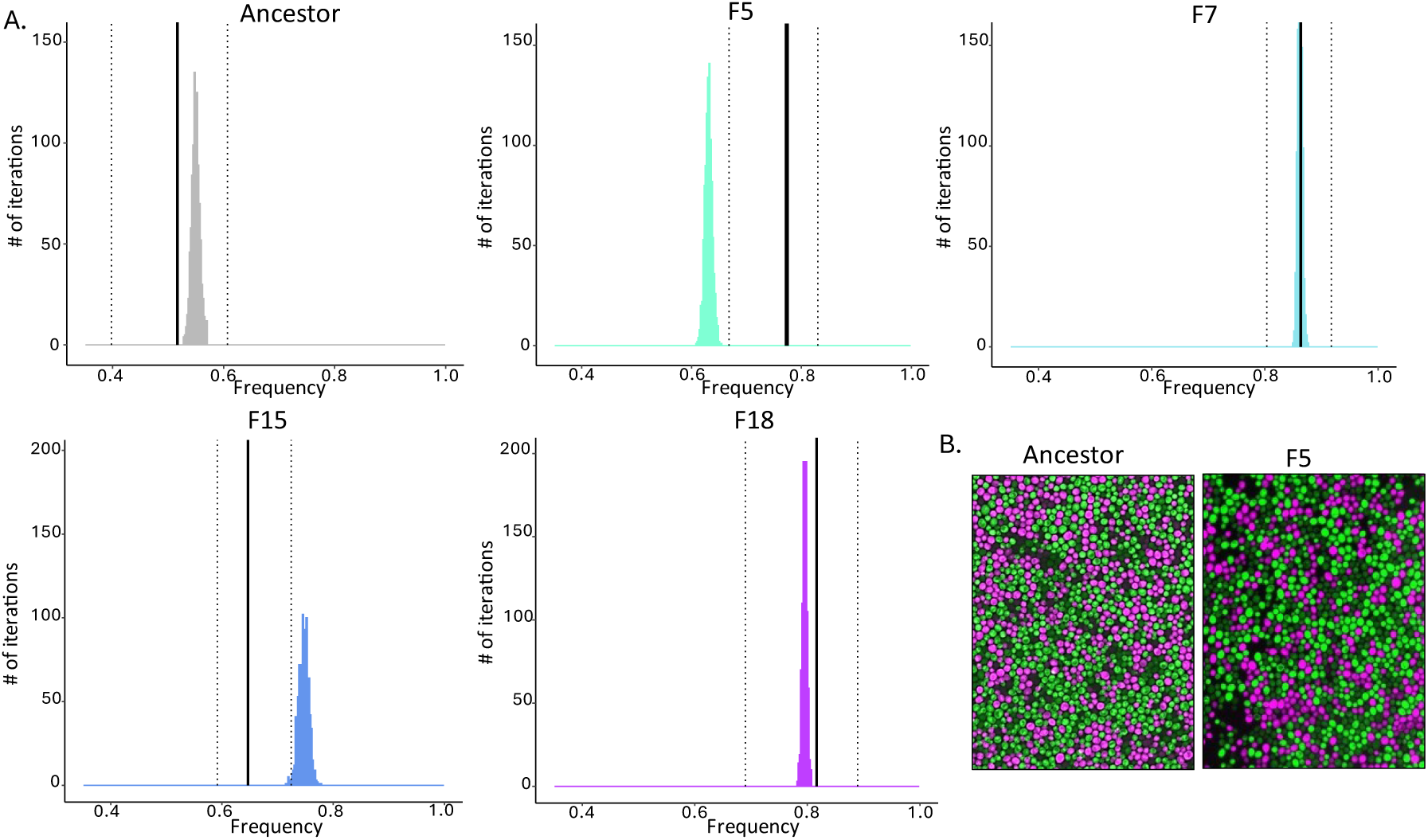
Spatial structure within flocs. A. Comparison of observed versus expected cell-cell interactions within flocs. Histograms show null distributions generated from 1000 simulations of randomly distributed cells, while vertical lines show the actual observed data: solid lines indicate mean observed interaction frequencies between evolved cells, and dotted lines show their 95% confidence intervals. F5 showed significant positive spatial structure, with observed interactions 22.5% higher than expected and all simulated frequencies falling outside the observed confidence interval. F15 showed significant negative spatial structure, with observed interactions 15% lower than expected and 99% of simulated frequencies above the observed confidence interval. Ancestor, F7, and F18 showed no spatial structure, with simulated frequencies falling within observed confidence intervals. B. Representative microscopy images of floc isolates flattened into a 2D distribution. Ancestor controls were made by co-culturing GFP- and RFP-labeled cells.

For the F5 lineage, the mean frequency of observed interactions between evolved cells was 22.5% higher than expected at random, and all simulated interaction frequencies fell outside of the 95% confidence intervals of the observed data (Figure 5A). This spatial structure, combined with our earlier finding that the F5 *FLO1* gene alone was sufficient for positive assortment, provides compelling evidence for the evolution of kin recognition in this lineage.

In contrast, lineage F15 showed an unexpected pattern: evolved cells had significantly fewer interactions than predicted at random. The null distribution exhibited a 15% higher mean frequency of interactions between evolved cells, with 99% of simulated values falling outside the 95% confidence interval of the observed data (Figure 5A). This suggests evolved F15 cells maintained greater spacing between one another than expected by chance. The F15 lineage evolved a mutation replacing a serine residue with a bulky, hydrophobic phenylalanine adjacent to the binding site (Figure 2). When removed from its evolved cellular context, this mutated *FLO1* allele lost function (Figure 3), suggesting the F15 *FLO1* gene evolved in concert with other cellular components. These changes appear to have fundamentally altered cell-cell interactions, reducing compatibility with unrelated cells. However, future work will be necessary to fully understand the mechanism of assortment in this lineage. The ancestor, F7, and F18 lineages showed no evidence of spatial structure - their null distributions fell completely within the 95% confidence intervals of the observed data (Figure 5A). Thus, it appears that these strains evolved a kin-agnostic increase in adhesiveness.

## Discussion

We tested whether prolonged selection for large multicellular size favored the evolution of positive assortment in aggregative *S. cerevisiae*. Evolved yeast were co-cultured with the ancestor, and we calculated assortment by measuring the relative frequency of evolved cells in flocs normalized by their overall frequency in the population. We observed the evolution of positive assortment for 13 out of 19 evolved lineages. During experimental evolution, four lineages convergently evolved point mutations in the *FLO1* gene (Pentz *et al*. 2023), which plays a role in mediating cell-cell aggregation. Leveraging these *FLO1*-mutant lineages, we examined potential mechanisms underpinning the evolution of positive assortment using genetic engineering, competitive inhibition assays, and analysis of spatial structure within flocs. We observed multiple potential independent evolutionary pathways resulting in positive assortment. A single lineage (F5) exhibited compelling evidence for differential recognition and adhesion between cells driven by changes to the *FLO1* gene, consistent with kin recognition (Hamilton 1964, 1970; McDonald et al. 2017). However, the three remaining *FLO1*-mutant lineages did not show evidence of kin recognition, and assortment appeared to result from traits that generally improved the flocculation efficiency of cells but did not result from genetic specificity. Moreover, lineages exhibiting positive assortment did not gain a fitness advantage during settling selection, suggesting that positive assortment evolved as an indirect effect of changes to cell-level traits that improved flocculation, but cells are not under social selection.

The lack of adaptive benefit generally exhibited across assortative strains was unexpected - evolved cells settled an average of 12-fold faster than the ancestor in isolation. This suggested that cells capable of preferentially aggregating with similarly fast-settling clonemates should gain substantial fitness advantages. In part, the modest levels of assortment we observed help explain this apparent paradox. Assortment values averaged 0.176 and ranged from 0.068 to a high value of just 0.328 (Table S1), indicating that ancestral cells could readily infiltrate even the most assortative flocs. However, while these assortment values appear modest, they represent substantial evolutionary changes from the ancestor. For instance, lineage F4’s assortment of 0.068 marked a 2.86-fold increase, and lineage F16’s value of 0.328 represented a 13.8-fold change. These shifts suggest the potential for further increases in assortment through continued selection. As cells evolve adaptations that enhance flocculation efficiency, their increased survival and reproduction under size-based selection should lead to their growing representation both in the population and within individual flocs. This increasing assortment, though apparently arising as a byproduct of selection on flocculation efficiency, may prove crucial for the evolution of other multicellular traits. While not initially adaptive for settling speed, increased assortment could facilitate the evolution of collective-level traits that impose individual costs or require coordinated behavior across cells. Thus, these early increases in assortment may provide an important foundation for future multicellular adaptations.

The pathway by which positive assortment evolves may underpin opportunities for multicellular collectives to become new Darwinian individuals (Bourrat 2019; Godfrey-Smith 2019). We observed the evolution of positive assortment through at least two independent pathways. For example, our analyses of the F5 lineage suggests the evolution of positive assortment through the pathway of genetic recognition: when we placed the F5 FLO1 gene (containing mutation N84 in a predicted ‘hinge’ near the binding pocket of the protein, Figure 2) into the ancestral genomic background, these ANC+FLO1_evo_ cells maintained similar assortment levels as F5 cells. The F5 lineage also exhibited spatial structure within flocs, with cells preferentially binding to clonemates within flocs. Importantly, F5 flocs showed no increased resistance to competitive inhibition by exogenous mannose, indicating that their positive assortment did not simply result from enhanced cellular adhesion. Together, these results suggest that F5 cells evolved a kin recognition system mediated by their shared *FLO1* alleles.

In contrast, positive assortment can arise through indiscriminate aggregation, where cells become generally more adhesive regardless of partner genotype (Wolf et al. 1999; Smukalla et al. 2008). Such cells are more likely to form genetically chimeric collectives, especially when multiple genotypes evolve similar adhesive properties. This pathway may limit the efficacy of selection on multicellular traits by impeding a connection between collective phenotype and genotype. We observed this pattern in the remaining *FLO1*-mutant lineages (F7, F15, F18). Changes to *FLO1* alone did not explain their positive assortment, they lacked spatial structure within flocs, and they showed significant resistance to exogenous mannose, suggesting their assortment stemmed from generally enhanced adhesion. However, for F15, the disrupted function of *FLO1* when removed from its evolved cellular context suggests possible interactions between *FLO1* and other cellular components.

In general, evolutionary transitions of aggregative multicellular organisms face significant challenges and eco-evolutionary tradeoffs. Our study revealed a striking pattern: while 13 of 19 evolved lineages developed positive assortment, it did not generally offer an adaptive benefit - a surprising result given that these lineages settled an average of 12-fold faster than their ancestor when grown in isolation. Theory predicts that preferential assortment should provide substantial evolutionary benefits by reducing within-group conflict and increasing the efficacy of selection on collective-level traits (Price 1970; Fisher et al. 2013; Pentz et al. 2020). The apparent lack of such benefits suggests competing selective pressures. Consistent with this, while not statistically significant, we observe a negative correlation (*Pearson’s r* = -0.299, *95% CI* [-0.66, 0.18]) between assortment and competitive settling success, indicating assortment may even carry weak costs. Only one *FLO1*-mutant lineage showed evidence of kin recognition-based assortment. While this selective partner choice could provide long-term evolutionary advantages by preventing chimeric aggregates, it may carry immediate ecological costs through slower aggregation (Pentz et al. 2020). The remaining lineages we studied appear to have evolved positive assortment through mechanisms that enhanced general adhesion. This strategy may offer short-term ecological advantages - cells that quickly and indiscriminately join aggregates can outcompete more selective cells, but may limit their capacity for groups of cells to serve as units of selection, gaining multicellular adaptations in response to selection acting on the traits of groups.

Whether a positively assorted lineage gains an adaptive benefit may depend on the evolutionary pathway by which assortment evolves. In this study, we observed at least two independent pathways to assortment, but we only evaluated mechanisms for the four lineages that evolved mutations in *FLO1*. Future work should aim to thoroughly identify specific mechanisms underlying assortment and test whether the costs of assortment vary across distinct evolutionary pathways. Moreover, future work should examine whether pleiotropic constraints associated with selection for rapid growth limit the benefit of assortment for settling success. We hypothesize that eco-evolutionary tradeoffs associated with distinct evolutionary pathways as the most likely explanation for the limited adaptive benefit associated with positive assortment.

This study highlights fundamental constraints on the evolution of aggregative multicellularity. Across dozens of independent transitions to multicellularity, no organisms that form multicellular groups through aggregation have evolved ‘complex’ multicellular morphologies (Knoll 2011; Brunet and King 2017). Our findings offer insight into why this might be: despite strong selection for rapid settling that drove a 12-fold increase in settling speed across 20 replicate populations of flocculating yeast, positive assortment provided no adaptive benefit when it evolved. The lack of a benefit from assortment, coupled with generally low assortment values, suggests that genetic associations between cells arose indirectly as a result of selection for partner-agnostic adhesion, rather than through as a result of inter-group competition during settling selection. These results demonstrate significant constraints on the evolutionary transition to multicellular Darwinian Individuality in our aggregative system, even under conditions entailing strong between-group selection. The assortment we observed might nonetheless represent early stages of this evolutionary transition: multicellular adaptation could emerge gradually as selection on cell-level traits indirectly shapes collective properties, eventually providing a foundation which allows selection on collective-level traits to drive collective-level adaptation

## Methods

### Yeast strains, experimental evolution, and media

Strains used in this study are described in Supplemental Table 2. The construction of the ancestor to the experimentally evolved strains used in this study is described in Pentz *et al*., 2020. The ancestral genotype was created from a homozygous diploid unicellular background (*Saccharomyces cerevisiae* strain Y55, accession JRIF00000000). Yeast were designed to flocculate by replacing the *URA3* open reading frame (ORF) with the KAN-GAL1p::FLO1 cassette (Smukalla *et al*., 2008). This ancestral genotype was used to initiate experimental evolution of twenty isogenic, replicate populations. Experimental evolution of these replicate populations is explained in detail in Pentz *et al*., 2023. Briefly, each replicate population was seeded into 10mL of YPGal+Dex [1.8% galactose (w/v), 0.2% dextrose (w/v), 2% peptone (w/v), 1% yeast extract (w/v)], and replicate populations were then subjected to daily growth and settling selection. Cells were grown for 24 hours before 1.5mL of each overnight culture was removed and placed into a 2mL microcentrifuge tube. Settling selection was performed by allowing cells to settle to the bottom of the tube for five minutes, and the top 1.4mL were removed and discarded. The remaining 100uL of culture was transferred to 10mL of fresh YPGal + Dextrose for the next round of selection. Whole populations were cryogenically stored at -80 °C every seven days. After 24 weeks of evolution (168 rounds of selection), a single representative strain isolate was selected from each population. These representative strains (strains F1-F20) were used in this study. Four of these representative strains evolved nonsynonymous mutations in the *FLO1* gene (strains F5, F7, F15, and F18), and these *FLO1*-mutants were used to identify mechanisms underpinning positive assortment (Figure 2).

### Genetic Engineering of Experimentally Evolved Strains

To characterize strains with fluorescence microscopy and flow cytometry, we created constitutive GFP and RFP tagged versions of each of the 20 evolved flocculating yeast isolates described in this study (Supplemental Table 2). Fluorescent markers were inserted at the *LYS2* locus, interrupting one copy of *LYS2 a*nd leaving the other intact. This was done using PCR to amplify GFP or RFP (along with the NAT drug resistance marker) from plasmids pYM25 and pYM42, respectively, with primers whose overhangs are homologous to the *LYS2* locus. These PCR products were separately transformed into each strain F1-F20 and selected for on YPD plates containing nourseothricin.

We additionally created strains containing *FLO1* alleles from evolved strains that had obtained a mutation in the *FLO1* reading frame (F5, F7, F15, and F18), in a genomic background that otherwise matched the ancestor (referred to as ANC+FLO1_evo_ strains; Supplemental Table 1). This was done using PCR to amplify the region from evolved strains containing the KanMX marker, *FLO1* gene, and some of the surrounding *URA3* locus. These PCR products were then transformed into Y55, the same strain that had been transformed with the *FLO1* gene to create the flocculating ancestor, with transformants selected on YPD with G418. Sporulation media (2% lithium acetate) was used to induce sporulation, and tetrads were dissected to obtain strains homozygous for the evolved *FLO1* allele via self-diploidization. These strains were then tagged with GFP as described above.

### Measurements of Floc Assortment

To begin assortment analysis, GFP-labeled representative strains from each replicate population (strains F1-F19) were revived by streaking onto YPD agar plates. The ancestor was separately labeled with two fluorescent markers (GFP and RFP), and we revived the unlabeled, RFP- and GFP-labeled ancestors by streaking onto YPD agar plates. All revived strains were grown at 30 °C for 48 hours. After 48 hours, overnight liquid cultures were prepared by swiping cells from the densest region of growth on the streak plates (limiting the probability of selecting a spontaneous mutant using a single CFU), and cells were inoculated into 10mL YPGal+Dex. Cultures were grown for 24 hours at 30 °C with shaking at 250 rpm. Overnight cultures were vortexed, and 1mL was removed from each sample. YPGal+Dex was removed by centrifuging and cells were deflocculated by mixing in 100mM EDTA. While remaining in EDTA, co-cultures were prepared by mixing each GFP-labeled evolved strain with the RFP-labeled ancestor in equal volumetric ratios (200uL of each strain = total volume of 400uL). A control was also prepared by mixing the GFP-labeled ancestor with the RFP-labeled ancestor (200uL of each). Mixed cultures were then centrifuged at 6000 rpm, EDTA was removed, and the co-cultured cells were resuspended in 400uL YPGal+Dex. Overnight co-cultures were prepared by placing 200uL of the cell mixtures into 10mL YPGal+Dex and incubated for 24 hours at 30 °C with shaking at 250 rpm. We also performed the protocol described here for unlabeled, RFP-, and GFP-labeled ancestor cells that were not co-cultured with another strain and served as controls during flow cytometry.

After 24 hours, co-cultures and controls were removed from the incubator one at a time to preserve floc integrity (flocs can flatten when sitting in stationary cultures). To prepare “floc” samples, flocs were carefully removed from each culture, limiting the uptake of unicellular cells suspended in culture media. If excess culture media was incidentally acquired with flocs, it was removed after allowing the flocs to settle in microcentrifuge tubes. Cells were then deflocculated by mixing in 100mM EDTA. Deflocculated cells were centrifuged at 6000 rpm, EDTA was removed, and the remaining cells were resuspended in 200uL sterile DI water. Samples for flow cytometry were prepared by suspending 50uL of the prepared floc cells into 1.5mL sterile DI water. “Population” samples were prepared to control for the overall frequency of evolved cells in the population (*i.e*., the total frequency of evolved cells in flocs and as unicells). To prepare “population” samples, overnight cultures were vortexed, and 1mL was removed from each culture. Cells were deflocculated by mixing in 100mM EDTA. Deflocculated cells were then centrifuged at 6000 rpm, the EDTA was removed, and cells were resuspended in 500uL sterile DI water. Samples for flow cytometry were prepared by placing 50uL of the prepared population cells into 2mL sterile DI water. We also performed the protocol described here for the singly cultured unlabeled, RFP-, and GFP-labeled ancestor controls. The relative frequencies of RFP-and GFP-labeled cells in each culture were quantified using a BD FACS Melody analyzer. Gates were set using the unlabeled and single-color ancestral controls. Assortment values were calculated according to Eq. 1.

We then repeated the assays described above and measured assortment of ANC+FLO1_evo_ cells when they were co-cultured with the ancestor. We also used these lineages to test whether positive assortment depended on recognition of both a partner’s FLO1 protein and its cell wall context. To do this, we co-cultured ANC+FLO1_evo_ cells with evolved cells sharing the same *FLO1* allele (e.g., F5 ANC+FLO1_evo_ with F5). Both of these assortment assays followed the same protocol as described above and assortment values were calculated according to Eq. 1. Each assortment assay was independently replicated four times.

### Competitive Settling Success

Competitive settling success was measured and the selection rate constant (Lenski et al. 1991) for each lineage was calculated as described in Pentz *et al*., 2023. Briefly, GFP-labeled evolved cells were co-cultured with RFP-labeled ancestor cells, as described above. Co-cultured cells were then subjected to a single round of growth and settling selection. The relative frequencies of GFP-labeled evolved and RFP-labeled ancestor cells were assessed using fluorescent microscopy. The selective advantage of evolved cells was measured as the relative fitness of evolved cells during settling selection. In this study, we leveraged this data collected by Pentz *et al*., 2023 and performed a linear regression to assess whether the competitive success during settling selection is a function of assortment value.

### Protein structure simulation and visualization

Structures of the N-terminus of FLO1p were simulated using AlphaFold 3, via alphafoldserver.com (Abramson et al. 2024). Residues 1-277 of the wild type and mutant (S51F, N84D, K194T, and P201R) *FLO1* gene sequences without any ligands were used as input. Residues 26-269 were superimposed for structural comparison in order to exclude disordered signal peptides and amyloid regions, using Mol* Viewer (Berman et al. 2000; Sehnal et al. 2021). PDB structure 4LHN (Goossens et al. 2015; Lelasi and Willaert 2023) of the N-terminal domain of FLO1 was also visualized via Mol* viewer.

### Mannose Competitive Inhibition Assays

Because we did not observe strong evidence that changes to the *FLO1* gene alone underpinned the positive assortment of the evolved strains, we predicted that changes to other cell wall components contributed to assortment. Cellular adhesion occurs through recognition of the FLO1 protein but is mediated by interactions between mannose oligosaccharides that attach to a partner’s cell wall. We predicted that changes to mannose production or expression allowed evolved cells to form stronger bonds. To test this prediction, we performed mannose competitive inhibition assays. To prepare mannose media, we prepared YPGal+Dex as described above. We then added mannose to the YPGal+Dex media in small (100mL) batches that contained mannose at final concentrations of 0%, 1%, 2.5%, 5%, and 7%. This approach ensured that all samples tested in a single replicate assay received the same media but limited the waste of costly mannose that may go unused by preparing large batches.

Each replicate assay began by reviving the ancestor and *FLO1*-mutant strains from glycerol stocks by streaking onto YPD agar plates. Revived strains were incubated at 30 °C for 48 hours. Liquid overnight cultures were prepared by swiping cells from the densest region of growth on streak plates and inoculated into 10mL YPGal+Dex. Cells were grown overnight at 30 °C with shaking at 250 rpm. After 24 hours, each culture was vortexed, and 1mL of each culture was transferred to fresh YPGal+Dex. Cultures were then grown for an additional 24 hours at 30 °C with shaking at 250 rpm. Each overnight culture was then vortexed, and 1mL of each culture was transferred to 10mL of YPGal+Dex+Man at each of the 0%, 1%, 2.5%, 5%, and 7% concentrations. Cultures were incubated for 24 hours at 30 °C with shaking at 250 rpm. A control culture was also prepared by inoculating the ancestor into YPD (lacking galactose), so that flocculation was not induced.

The cell cultures prepared above were used to perform flocculation efficiency assays. Cultures were removed from the incubator in random order one at a time. Samples were vortexed, and we transferred 3mL of each sample into three transparent cuvettes (3mL/cuvette x 3 cuvettes = 9mL analyzed per culture). Cuvettes were placed on a backlit stage, making flocs visible in liquid media. Cultures were synchronously mixed using an automated multi-channel pipette, and we captured a video of the flocs suspended in solution. High resolution still images were captured from each video, and we assessed flocculation efficiency using gray-level co-occurrence matrices (GCLM) to measure pixel dissimilarity across each image. Specifically, dissimilarity measurements were performed by measuring the difference in pixel values between each pixel and all neighboring pixels. The sum of the absolute difference in pixel values over the entire image was calculated to determine the heterogeneity in pixels across the image. High dissimilarity values indicate high heterogeneity, resulting from a large amount of cell clumping. Low dissimilarity indicates cultures are homogeneous with a low amount of cell clumping. We determined the dissimilarity value, reported as the “flocculation efficiency,” for each culture by using the mean value across the three technical replicate images. Each mannose competitive inhibition assay was independently replicated four times.

### Within Floc Spatial Structure Analysis

To perform spatial structure analysis, co-cultures of GFP-labeled evolved cells and RFP-labeled ancestor cells were prepared as described for the measurements of floc assortment. Overnight co-cultures were then removed from the incubator one at a time. If the flocs were small, a single floc was removed, placed on a microscope slide, and gently flattened using a cover slip until cells formed a two-dimensional distribution without overlap. If flocs were large, we isolated a portion of the floc, which was similarly flattened. We captured images of each flatted floc using a Nikon Eclipse Ti-2 inverted microscope at 40x magnification. Images were cropped manually cropped to remove blurry regions and blank space then processed using ImageJ. The channels of each image were split, images were converted to grayscale, the threshold of each image was set using the default and auto settings, regions of interest (ROIs) were detected by analyzing particles with a circularity of at least 0.5, and cell separation was improved using the watershed function. ROI detection was performed using an ImageJ macro, but each image was manually evaluated, and manual corrections were made as necessary. We recorded the coordinates for the center of each ROI, which were then used for spatial structure analysis.

We tested for spatial structure by measuring the frequency of interactions each individual evolved cell shared with another evolved cell (GFP-GFP) versus with the ancestor (GFP-RFP). Interaction frequencies were determined by measuring the interaction types within a 4uM radius. We chose this distance because measurements were made starting at the center of the focal cell, and the radius of a single cell is approximately 2.5uM. We also performed measurements at several increasingly large radii (6uM, 8uM, and 10uM), but we did not observe interesting patterns of spatial structure. This is not surprising considering using larger radii would capture “interactions” between non-adjacent cells.

We determined whether the observed frequency of interactions between cell types was different than expected at random, by comparing observed interactions frequencies to a simulated null distribution. To do this, we extracted the image dimensions from each sample image (images had been individually cropped to remove blurry cells and blank space). We then extracted the total number of GFP (evolved) and RFP (ancestor) cells present in each image. We used the mean image dimensions and numbers of GFP and RFP cells across the three replicates for each lineage to simulate the random distribution of GFP and RFP cells. A null distribution of interaction frequencies was created by recording the frequency of GFP-GFP interactions (or interactions between evolved cells) over 1000 simulation iterations. We compared the null distribution to the observed data and assessed the proportion of times the simulated frequency of interactions between evolved cells (GFP-GFP interactions) fell outside of the 95% confidence intervals of the observed interaction frequencies for each lineage. Simulations of expected cell-type interactions were performed using Python version 3.11.4.

### Statistics

To analyze whether evolved strains exhibited assortment, we used repeated t-tests with a Benjamini-Hochberg correction to test for differences from the ancestor. The adaptive benefit of positive assortment was assessed using a linear regression to determine whether competitive fitness during settling selection was a function of assortment value. Linear regression was performed using the R package ‘lme4’ (Bates et al. 2015). To investigate how the flocculation efficiency of the evolved lineages differed from the ancestor in response to increasing mannose concentrations, we fit a linear regression with an interaction term between lineage and mannose concentration (treated as a continuous variable).

Flocculation efficiencies of each evolved lineage were normalized by the ancestor. We normalized the data by subtracting the ancestral value from each assay replicate from the flocculation efficiency values of each evolved lineage, making ancestral flocculation efficiencies equal to zero. We then compared the slope of the regression for each evolved lineage to that of the ancestor (m = 0) to assess overall different trends in flocculation efficiency as mannose concentration increased. Differences in flocculation efficiency relative to the ancestor at 0% mannose, were determined by comparing the intercepts of the evolved lineages to that of the ancestor. Differences in flocculation efficiency at each specific mannose concentration were analyzed by deriving estimated marginal means from the model, and predicting the pairwise relationships between lineages using the R package ‘emmeans’ (Lenth 2022). Post-hoc comparisons between lineages at each concentration were performed using the ‘contrast’ function, and we applied a Bonferroni correction to control for multiple comparisons. We used the R packages ‘dplyr’ (Wickham H, François R, Henry L 2022) to prepare data and ‘ggplot2’ (Whickam 2016) for data visualization. All statistical analyses were performed in R version 4.2.2.

## Author contributions

K.S.S., K.A.M., and W.C.R. designed experiments and analyzed the data. K.A.M. and A.J.B. performed genetic engineering of yeast strains. Experiments were conducted by K.S.S. and C.B., and protein fold simulations were performed by A.J.B. K.S.S., K.A.M., and W.C.R. wrote the first draft of the manuscript, and all authors contributed to revisions. All authors approve of the final version of the manuscript.

## Acknowledgements

We would like to thank Jennifer Pentz for her comments and sharing of prior data. We would also like to thank members of the Ratcliff lab for their comments on the manuscript.

