## Supplemental Table 1 for "Multiple pathways to the evolution of positive assortment in aggregative multicellularity"

| **Table 1. Assortment valued and statistics for evolved lineages.** | | | | | |
| --- | --- | --- | --- | --- | --- |
| Lineage | FLO1 | Assortment (r) | df | t-statistic | p-value |
| F4 | Mutant | 0.068 | 16 | -2.64 | **0.025** |
| F5 | Mutant | 0.148 | 16 | -3.73 | **0.004** |
| F7 | Mutant | 0.033 | 16 | -0.71 | 0.51 |
| F15 | Mutant | 0.215 | 16 | -5.00 | **0.0004** |
| F18 | Mutant | 0.167 | 16 | -3.64 | **0.004** |
| F1 | Non-mutant | 0.114 | 16 | -2.92 | **0.015** |
| F2 | Non-mutant | 0.246 | 16 | -5.87 | **0.0001** |
| F3 | Non-mutant | -0.049 | 16 | 0.47 | 0.64 |
| F6 | Non-mutant | -0.061 | 16 | 1.00 | 0.39 |
| F8 | Non-mutant | 0.246 | 16 | -5.01 | **0.0004** |
| F9 | Non-mutant | -0.198 | 16 | 2.61 | **0.025** |
| F10 | Non-mutant | -0.066 | 16 | 0.79 | 0.49 |
| F11 | Non-mutant | 0.158 | 16 | -2.91 | 0.16 |
| F12 | Non-mutant | -0.076 | 16 | 1.13 | 0.35 |
| F13 | Non-mutant | 0.164 | 16 | -5.31 | **0.0003** |
| F14 | Non-mutant | 0.295 | 16 | -8.92 | **<0.0001** |
| F16 | Non-mutant | 0.328 | 16 | -9.84 | **<0.0001** |
| F17 | Non-mutant | 0.262 | 16 | -4.22 | **0.001** |
| F19 | Non-mutant | 0.144 | 16 | -4.58 | **0.0008** |
| Note: Assortment values were determined by co-culturing evolved lineages with the ancestor and measuring the frequency of evolved cells in flocs relative to their frequency in the population, according to Equation 1. Changes to assortment levels were statistically analyzed by comparing the assortment values of the evolved lineages to the assortment of the ancestor using repeated t-tests with a Benjamini-Hochberg correction. We observed a significant increase in positive assortment for 13/19 lineages. Two lineages exhibited a change from the ancestor toward negative assortment (indicated by downward facing arrow in front of p-value). Bolded p-values indicate a significant difference from ancestor. Positive assortment was observed for lineages where mutations evolved at the FLO1 gene (mutant) and in lineages where they did not (non-mutant). | | | | | |
